# Association theory: a new framework for analyzing social evolution

**DOI:** 10.1101/197632

**Authors:** Owen M. Gilbert

## Abstract

The dominant social-evolutionary paradigm implicitly equates social actions and behaviors causing associations by extrapolating from models of social actions to explain behaviors affecting association. This extrapolation occurs when models of helping behavior are applied to explain aggregation or fusion, and when models of discriminatory helping behavior are applied to explain discriminatory segregation or discriminatory rejection. Here, I outline an alternative theoretical approach that explicitly distinguishes a social action as a helping or harming behavior, and an association as the context for a social action. Based on this distinction, I define a list of terms that allows a classification of association phenomena and the conceptual framework necessary to explain their evolution. I apply the resulting theory, which I call “association theory,” to identify a series of steps common to major and minor transitions in social evolution. These steps include the evolution of association, the evolution of differential treatment, the evolution of association preference, and the evolution of genetic kin recognition. I explain how to measure the parameters of association theory and I apply the theory to test Hamilton’s rule. I evaluate the evidence for association theory, including how it resolves anomalies of a former paradigm. Finally, I discuss association theory’s assumptions, and I explain why it may become the dominant framework for analyzing social evolution.

## Introduction

The history of life is punctuated by a series of major evolutionary transitions in hierarchical complexity and ecological diversity (Buss 1987; Bourke 2011). In some of the most consequential transitions in evolution, the evolution of altruism led to division of labor, specialization of castes, and the rise of emergent unicellular, multicellular, and colonial organisms (Maynard Smith and Szathmáry 1995; West et al. 2015). The quality of being “organismal” suggests complex traits at new hierarchical levels, allowed by functional integration and mutual interdependence of subparts (e.g., organs, tissues, castes; Strassmann and Queller 2010; Korb and Heinze 2016). In a number of minor transitions, organisms evolved to associate and, in some cases cooperate, but organismal degrees of integration and emergent complexity at new levels did not originate (Grosberg and Strathmann 2007; Fisher et al. 2013).

Found along with major and minor transitions in social evolution are traits that promote kin population structure. Examples of behaviors that actively promote kin population structure are those that cause organisms to segregate or reject based on variable cues shared with close kin (Buss 1982; Grosberg 1988; Grafen 1990). Such kin recognition systems help maintain the genetic integrity and individuality of organisms, including those that develop clonally and later fuse (Buss 1982, 1987; Bourke 2011). Traits promoting kin population structure are also found in organisms that lack cooperative behavior, for example fish that behave aggressively toward nonkin but associate preferentially with kin (Brown and Brown 1993).

Although traits promoting kin population structure occur in both major and minor transitions, they are thought to have their most important effects on major transitions, for example by controlling the spread of socially-destructive social parasites or facilitating the evolution of intraspecific cooperation (Buss 1982, Buss 1987; Bourke 2011). It is often suggested that traits promoting kin population evolve for long-term effects, as allowed by models assuming that multiple traits evolve simultaneously (Powers et al. 2011; Ryan et al. 2016). More generally, traits affecting population structure, including those that are discriminatory, are often explained with models of cooperation (Hamilton 1964; Vehrencamp 1983; Queller 1985; Buss and Green 1985; Rousset and Roze 2007; Bourke 2011).

Extrapolating models of cooperation to explain association phenomena yields a simple theory (West et al. 2015), but it yields three anomalies. First, it does not explain why species lacking altruism often restrict associations to kin (West-Eberhard 1989). Second, it does not explain why organisms that fuse often suffer substantial costs of fusion (Rinkevich and Weissman 1992). Third, it does not explain why “genetic kin recognition” cues, used for discriminatory segregation and rejection, are often highly variable (Rousset and Roze 2007).

The object of the present work is to outline a new framework for analyzing social evolution that resolves these anomalies, and which opens up new avenues for empirical and theoretical research. Here, I begin by explaining the fundamental distinction between a “social action” and “association,” two elementary concepts that have long been confused in social theory. I then develop a new theory, which I call “association theory,” based on this fundamental distinction. I define a list of terms (table 1) and core principles that follow from this distinction. Based on this conceptual framework, I outline a sequence of evolutionary steps that logically follow from it, and which I suggest commonly characterize social-evolutionary transitions (table 2). I then show how to measure the parameters of association theory in an empirical context, and how these parameters can be used to test Hamilton’s rule. I evaluate the evidence for association theory’s key predictions and review some of its assumptions. Finally, I explain why association theory may replace the current dominant paradigm of social theory.

**Table. 1:** Glossary.

| Term | Definition |
| --- | --- |
| Aggregation | The coming together of organisms, in which no groups coming together, as may distinguished by clonal genotype, are themselves discrete organisms before coming together (in contrast to fusion) |
| Association | The context for a social action |
| Association preference | Differentially entering into or avoiding the context for a social action based on a cue; e.g., due to discriminatory segregation or discriminatory rejection after initial aggregation or fusion |
| Automatic effects of association | The passive or incidental effects of being in the context for a social action, not due to a social action |
| Chimera | An organism derived from fusion or aggregation of more than one clone |
| Conflict | A social interaction involving harming behavior |
| Cooperation | A social interaction involving helping behavior |
| Cue | A factor, such as a phenotype or context, which is utilized to trigger a discriminatory behavior |
| Differential treatment | Changing the form of treatment (e.g., help or harm) based on a cue; discriminatory social action |
| Discriminate | With respect to factor *X*, expressed contingently on cues correlated with *X* |
| Exploitation | Pure selfishness, or separately, the cheating of altruism |
| Fusion | The coming together of two or more formerly-discrete organisms to form a single, chimeric or clonal organism |
| Indiscriminate | Not expressed contingently on cues correlated with factor *X*, though possibly expressed based on cues uncorrelated with *X*, obligately, or probabilistically |
| Nepotism | Kin favoritism; e.g., by preferentially helping or avoiding harming kin |
| Net effect of social interaction | The sum of the effects of relevant social actions on actors and recipients; usually positive for cooperation and negative for conflict |
| Obligate | Always expressed or constitutive |
| Organism | A living entity characterized by functional integration and mutual interdependence of subparts, such as proteins, cells, organs, tissues, or castes, which give rise to emergent complexity of the entity's traits |
| Social action | A helping or harming behavior toward another individual (recipient) |
| Social interaction | One or more social actions by one or more associated individuals |
| Treatment | The manner of acting toward another in a specified way (e.g., help or harm) |

**Table. 2:** Association preference and major transitions.

| Major Transition | Association Preference | Examples | Reference |
| --- | --- | --- | --- |
| **Non-Life → Unicellular** | Cellular fusion-rejection | Cell-cell recognition and outer membrane exchange in *Myxococcus xanthus* | Pathak et al. 2013 |
| **Uninucleate → Multinucleate** | Multinuclear fusion-rejection | Somatic compatibility in ascomycete fungi and plasmodial slime molds | Buss 1982<br>Saupé 2000<br>Clark and Haskins 2012 |
| **Unicellular → Multicellular** | Cellular aggregation-segregation | Kin-discriminatory segregation in social amoebae and territorial exclusion of nonkin in social bacteria | Velicer and Vos 2009<br>Hirose et al. 2011<br>Gilbert et al. 2012 |
|  | Multicellular fusion-rejection | Somatic compatibility or histocompatibility in marine invertebrates | Buss 1982<br>Grosberg 1988<br>Gilbert 2015 |
| **Multicellular → Colonial** | Multi-cellular aggregation-segregation | Discriminatory settlement in marine invertebrates, and kin association preference in terrestrial arthropods and aquatic and terrestrial vertebrates | Fletcher and Michener 1987<br>Grosberg 1988 |
|  | Colonial fusion-rejection | Nestmate discrimination in colonial arthropods and vertebrates | Fletcher and Michener 1987<br>Gamboa 2004 |

**Note:** Listed are the major types of association preference found in four major transitions of social evolution, as well as examples of taxon-specific terminology used. Aggregation-segregation association preference is indirectly associated with major transitions, useful for comparative study.

## Foundations

### Social Action Versus Association

I define a “social action” as a helping or harming behavior, and an “association” as the context for a social action (similar to Whitehead’s [2008] distinction between an “interaction” and an “association”). Conceivably, the context for a social action could be any number of temporal, spatial, or communicative contexts. In most cases, the context for a social action is provided by physical proximity or union (Simon 1962). Consequently, I focus on aggregation and fusion as behaviors that allow association between discrete organisms or within fused organisms, respectively.

The distinction between “social action” and “association” is not as specialized and technical as it sounds. For example, we often come into close proximity with others, as we walk to various places and go about our daily lives. Being in close proximity with others conveys us the ability to either help others (e.g., donate a dollar to a street musician) or harm others (e.g., steal a wallet), should we choose to. Had we stayed at home and not ventured out, we may not have been associated with others in these ways. However, we all know the difference between “association” and “social action,” since we can tell the difference between merely passing a street musician and placing a dollar in her empty guitar case. Not knowing the difference might, moreover, be quite costly. We might think that simply sitting next to somebody on a bench in a dark alley, to share body heat on a cold night, means that we are “cooperating.” We might believe that, until the person pulls a gun and robs us.

Distinguishing “social action” and “association” has important consequences for social theory. First, it illuminates a distinction between the “automatic” effects of association, which stem from being in the context for a social action, from the effects of social actions themselves. Automatic effects of association arise strictly from coming into the of a social action. In biological contexts, examples of automatic benefits of association include advantages of increased feeding efficiency (Parrish and Keshet 1999; Koschwanez et al. 2011), mobility (Parrish and Keshet 1999), ability to tolerate stressors (Allee 1931; Krause and Ruxton 2002), ability to avoid predators (Kessin et al. 1996; Sword et al. 2005), sharing of body heat (Gilbert et al. 2010), inheritance of territories or shelters (Krause and Ruxton 2002), sharing of information (Vogel and Dussutour 2016), and interspecific competitive ability (Buss 1981). Examples of automatic costs of association include parasite transmission (Davis and Brown 1999), attraction of predators (Krause and Ruxton 2002), decreased feeding efficiency (Krause and Ruxton 2002), break up of coadapted gene complexes (De Boer 1995), and depletion of local resources (Alexander 1974; Krause and Ruxton 2002). Automatic effects have also been called “accidental (Allee 1931)” or “passive (Kokko et al. 2001).” I use the term “automatic,” however, because automatic benefits of association may provide the initial selective pressure for association (thus, not being “accidental” or “passive” with respect to the adaptive value of association).

### Differential Treatment Versus Association Preference

Distinguishing a “social action” and an “association” allows a distinction between forms of discriminatory behavior. I refer to discriminatory social actions, like discriminatory help and harm, as “differential treatment.” In contrast, I refer to discriminatory behaviors that determine whether individuals enter the contexts for social actions as “association preference (table 1)”.

Like the distinction between “social action” and “association,” the distinction between “differential treatment” and “association preference” is one we all commonly make. As we all know from everyday experience, there is an important difference between whether we associate with somebody, and how we treat somebody given that we associate (or how they treat us). If we are smart, we might restrict our associations to those with whom we will cooperate rather than conflict. However, the reason for this, in contrast to the distinction itself, is not so obvious—it requires a conceptual model. To build this model also requires first understanding the mechanisms of differential treatment and association preference.

I assume here that differential treatment functions by the preferential helping or avoiding harming those sharing a cue, while association preference functions as an indiscriminate fusion or aggregation (association) behavior, followed by discriminatory rejection or segregation (avoidance) behavior. As an example of association preference, social amoebae *Dictyostelium discoidem* aggregate initially based on an acrasin stimulus (cyclic AMP) shared by members of the same species (Raper 1984), and then clones may segregate out based on polymorphic *tgr* cell adhesion genes (fig. *1A*; Hirose et al. 2011). Likewise, colonial tunicates *Botryllus schlosseri* fuse based on a species-specific activator of fusion encoded by the gene *fester* (Nyholm et al. 2006), and reject based on polymorphic *fuhc* (fig. *1B*; Taketa and De Tomaso 2015). Aggregation-segregation and fusion-rejection association preference are found across the tree of life (fig. 2), and along with four major transitions in social evolution (table 2).

**Figure 1:**
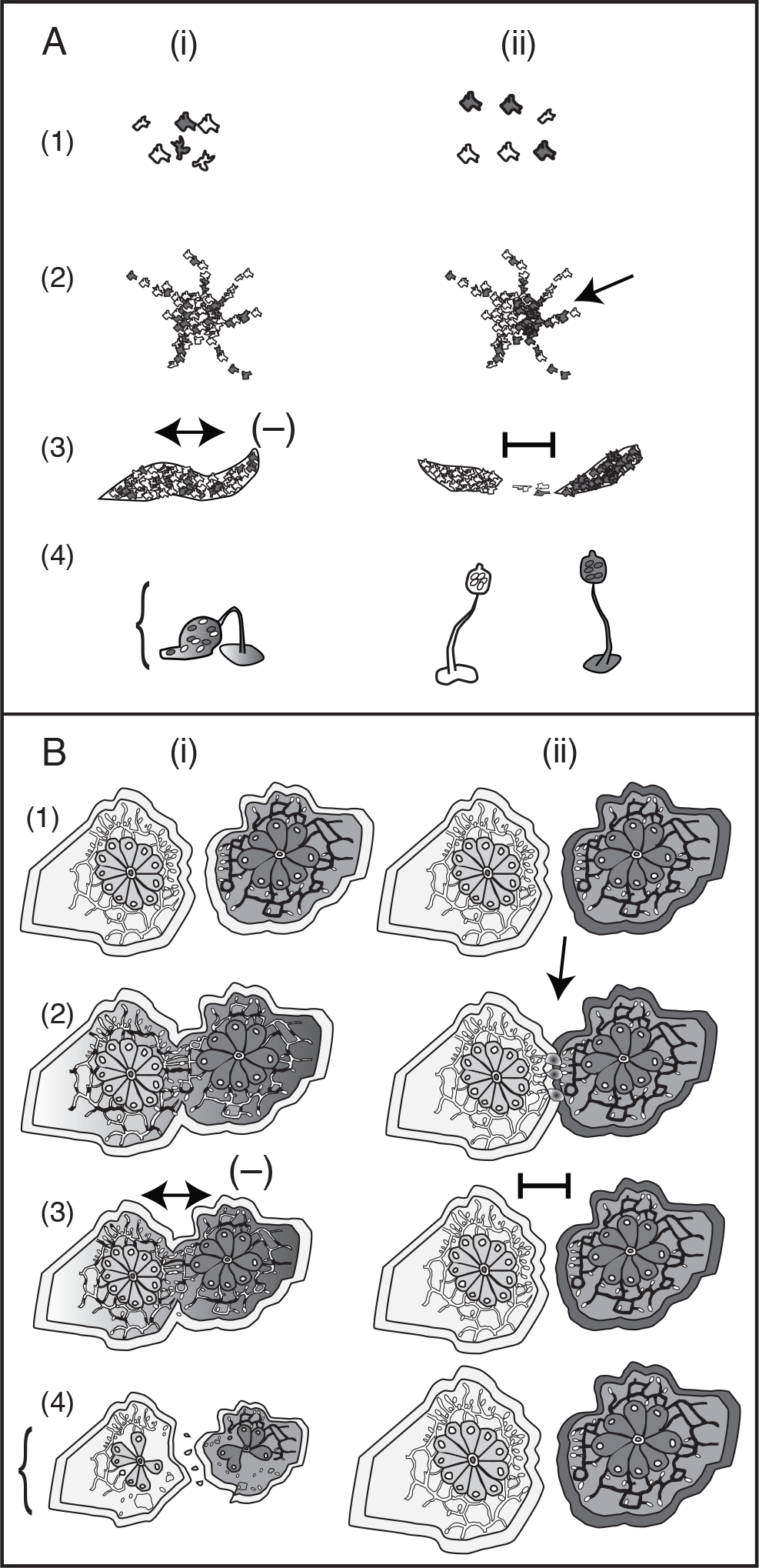
Examples of association, consequences differential treatment, and association preference. Encounters between conspecific clones, indicated by different shades, which lead to the potential for social actions. A, Social amoebae exhibit association as aggregation, which brings different clones together into the context for a social action. In column (i), clones differ at a gene that cues competitive programs for spore differentiation. In column (ii), different clones possess different functional alleles of cell adhesion tgr genes. Numbers (1) - (4) indicate sequential stages of events immediately before and following aggregation. In (i), aggregation leads to differential treatment and decreased stalk:spore ratios, resulting in unhealthy fruiting bodies (here, indicated by falling over). In (ii), aggregation leads to discriminatory segregation (arrow), which leads to fruiting bodies with correct stalk:spore ratios and thus normal relations with the environment. B, Marine invertebrates exhibit association as fusion, which brings different clones together into the context for a social action. In column (i), clones differ at a gene that cues competitive programs for germline parasitism, a form of differential treatment, indicated by the shade of the colony and vascular system. In column (ii), clones possess different functional alleles of the fuhc complex genes, indicated by the shade of the band around the colony. Numbers (1) - (4) indicate sequential stages of events immediately before and following fusion. In (i), fusion leads to mutual germline parasitism and shrinkage of colonies. In (ii), initial fusion leads immediately to discriminatory rejection (arrow), which prevents vascular fusion and maintains the health of both colonies.

**Figure 2:**
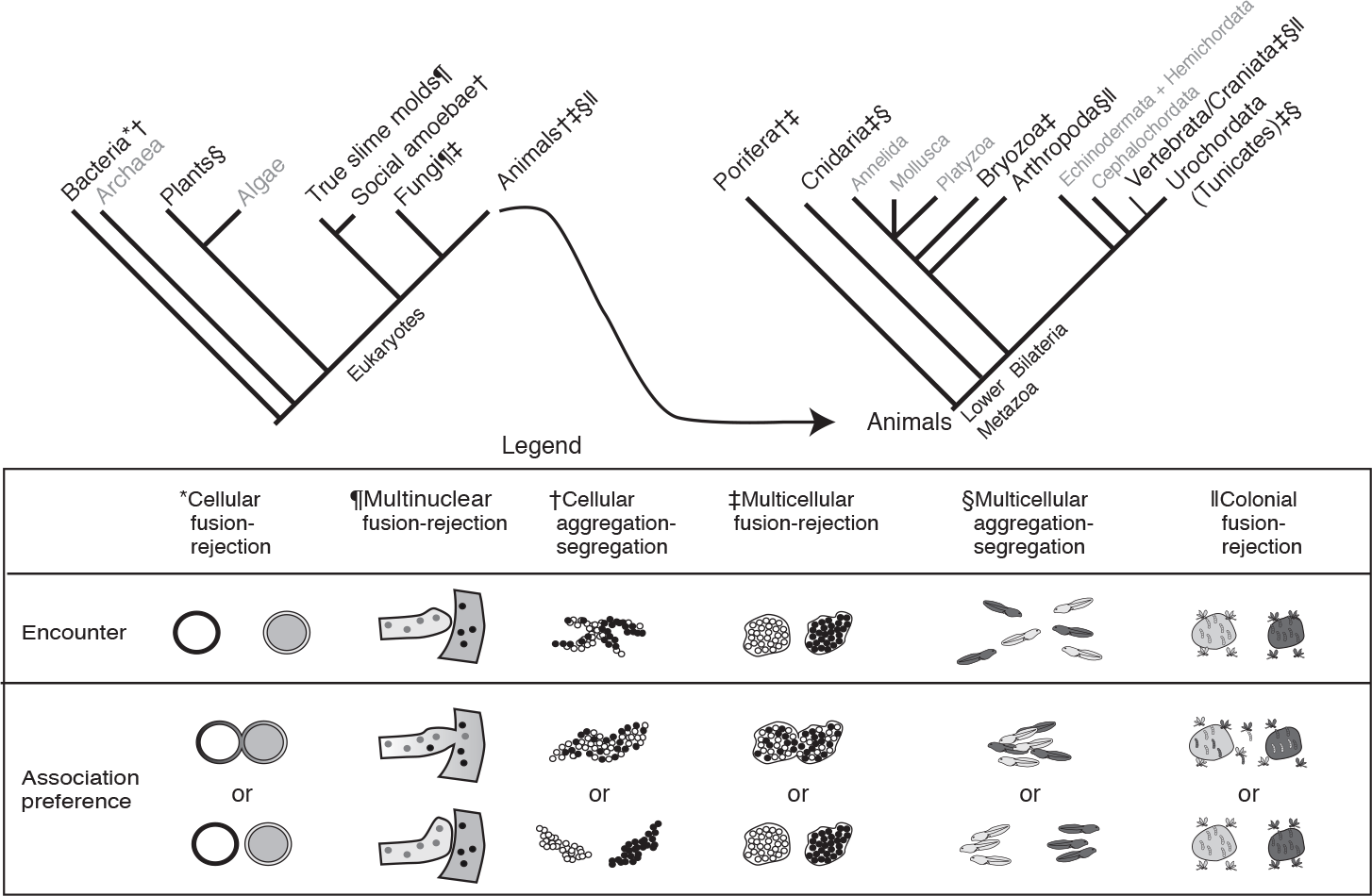
Phylogenetic distribution of association preference. The legend gives a description of the type of
association preference behavior found at each level of biological organization. The data suggest that association preference systems are a general theme of major transitions, but not ubiquitous. For example, some but not all kingdoms of complex multicellular organisms (Plants, Algae, Fungi, and Animals) exhibit multicellular fusion-rejection association preference (left-side phylogram; indicated by ‡); and some, but not all, animal phyla exhibit multicellular fusion-rejection association preference (right-side phylogram; also indicated by ‡). Cladograms are adapted from Rosengarten and Nicotra (2011). See text for details.

Many cases of aggregation and fusion can be captured with a pairwise model. Fusion is often pairwise, while aggregation is often pairwise with respect to genotype. For example, thousands of social amoebae might aggregate together to form a group, and yet such groups may usually be composed of one or two clones in nature (Gilbert et al. 2007). The question of association preference can thus be reduced to an encounter of two individuals, and whether they will associate. Two people approaching each other on a side walk have entered a context for a social action. A major conflict might nevertheless be avoided if one crosses the street. Likewise, both segregation and rejection “avoid” associations that initially begin with aggregation and fusion, respectively (fig. A1, available online).

Under my definitions (table 1), what primarily distinguishes “aggregation” from “fusion” is that the units associating, for example two clonal groups of amoebae, are not necessarily each “organismal.” Thus, the independent units could have developed themselves by “coming together” or “staying together” process. However, entities undergoing fusion will each have started life apart, and each being organismal, will have more likely been developed by a “staying together” process (Bourke 2011; Tarnita et al. 2013).

### Effects of Interactions

I assume that prior to associating, individuals cannot use cues to gauge asymmetries within associations, for example which particular individual will benefit more or less. For example, if two people are approaching each other on a side walk, they cannot gauge who is carrying a superior weapon until they are in the contexts in which weapons are normally drawn. Thus, I assume associations are “symmetric (Maynard Smith 1982).” However, I assume that cues might convey information about whether associations will be cooperative or conflicting. Thus, the two individuals approaching each other might discern traits (“cues”) that tip them off about the possibility of conflict or cooperation. Given that nonhuman organisms may often use cues indicative of genetic identity, I consider the consequences of helping and harming for the average effects of cooperative and conflicting social interactions, respectively (see table 1 and appendix A for my usage of “conflict”).

To conceptualize average or net effects of interactions, I break the standard terms of help and harm (Hamilton 1964) into components involving energy transfers. For help, I define *b* as the benefit to the recipient and *c* as the cost to the actor. For harm, I define *b* as the cost of harm to the recipient and *cș* as the benefit of harm to the actor. In terms of Hamilton’s rule, this means *b’* functions as the benefit of harm-avoidance to the recipient and *c’* as the cost of harm-avoidance to the actor (analogous to the *b* and *c* terms for help). I further break these terms into components, such that *b* = *e* + *e*_V_, and *c* = *e* + *e*_T_, where *e* is the value of the resource for the actor, *e*_V_ is the extra value of the resource for the recipient, and *e*_T_ is the cost of transferring the resource for the actor. Likewise, 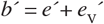, and 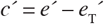, where *e’* is the value of the resource for the actor, 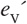’ is the extra value of the resource for the recipient, and 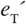 is the cost of transferring the resource for the actor. This reformulation of terms suggests cooperation will carry a net benefit, *B* = *b* – *c*, where *e*_V_ – *e*_T_ > 0, and that conflict will carry a net cost, *C* = *b′* – *c>′*, where 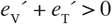 (fig. 3).

**Figure 3:**
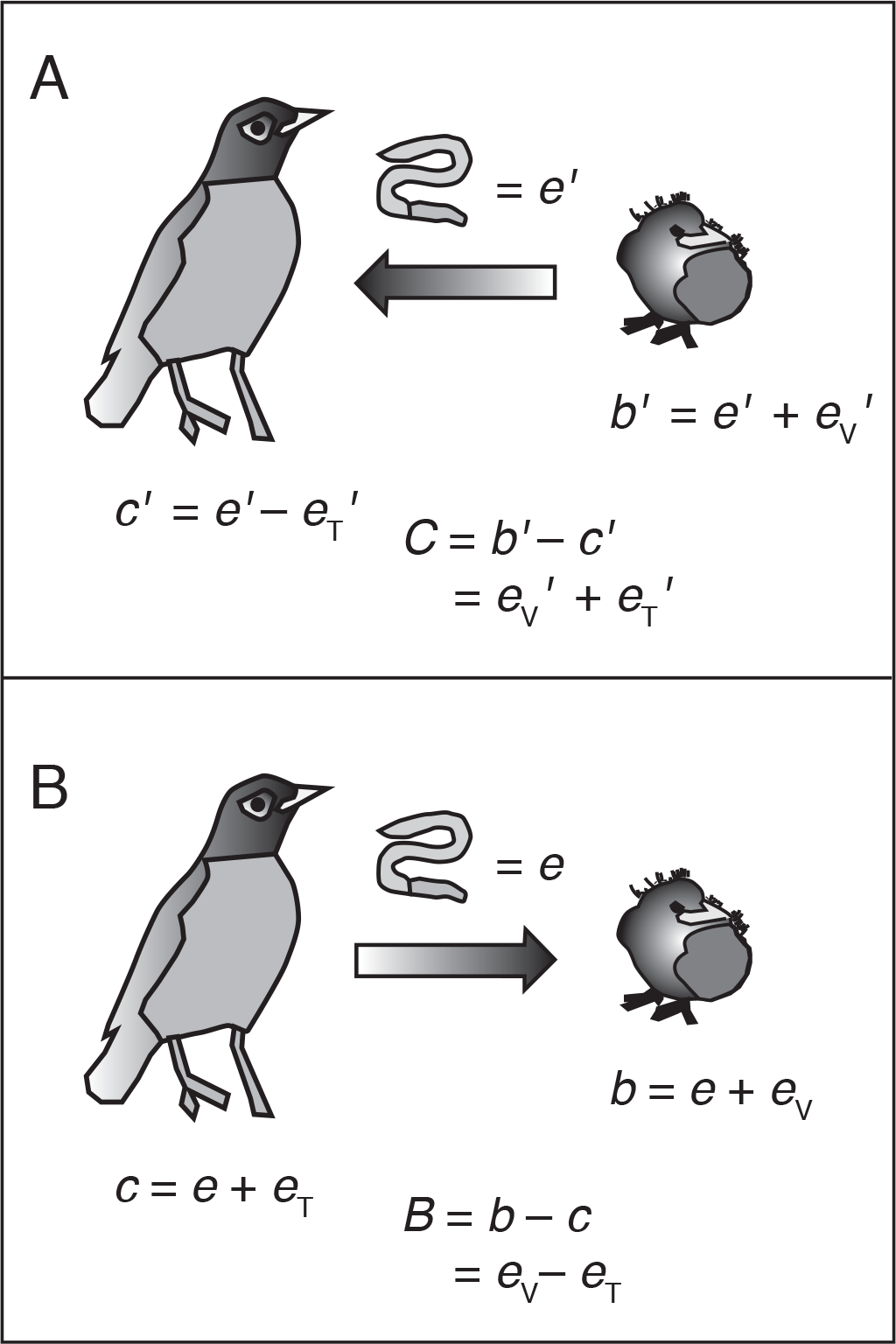
The net effects of interactions. The net effects of interactions. A, Conflict carries a net cost where the energy spent in transferring the resource for the actor plus the extra value of the resource to the recipient is positive 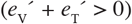. B, Cooperation carries a net benefit where the energetic cost of transferring a resource is less than the extra value of the resource to the recipient (*e*_T_ < *e*_V_). Here, the resource is a worm transferred between an adult and baby bird. See text for details.

This formulation of terms provides three insights. First, it suggests conflict will carry a net cost either where resources are more valuable to recipients, who are robbed, or where there is a sufficiently large cost of transferring the resources for actors (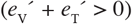). Situations where 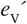 is likely to be positive where actors are stronger than recipients of harm, in which cases recipients of harm may suffer greater costs than the actors gain (e.g., a baby bird might be harmed more by losing a worm, than an adult gains by robbing the baby; fig. 3A). Second, it suggests cooperation will yield a net benefit where the extra value of the resource for the recipient is greater than the cost of transferring the resource for the actor (*e*_V_ > *e*_T_). *e*_V_ is likely to be positive are where an actor has superior skills in procuring or manufacturing a resource, as with division of labor or specialization (Maynard Smith and Szathmáry 1995; Bourke 2011). The same asymmetry of resource value can arise also where the actor is stronger, such that the same resource benefits the recipient more (fig. 3B). Third, where the resource being robbed *e′* is more valuable, more energy can be spent in robbing the resource. Consequently, the net cost of conflict can be larger where *e′* is greater (fig. 4). Biological situations where the greatest resources are at stake include cases where germlines are parasitized (Buss 1982), bodies cannibalized (Pfennig 1997), and nests or territories usurped (Field 1992; Buschinger 2009).

**Figure 4:**
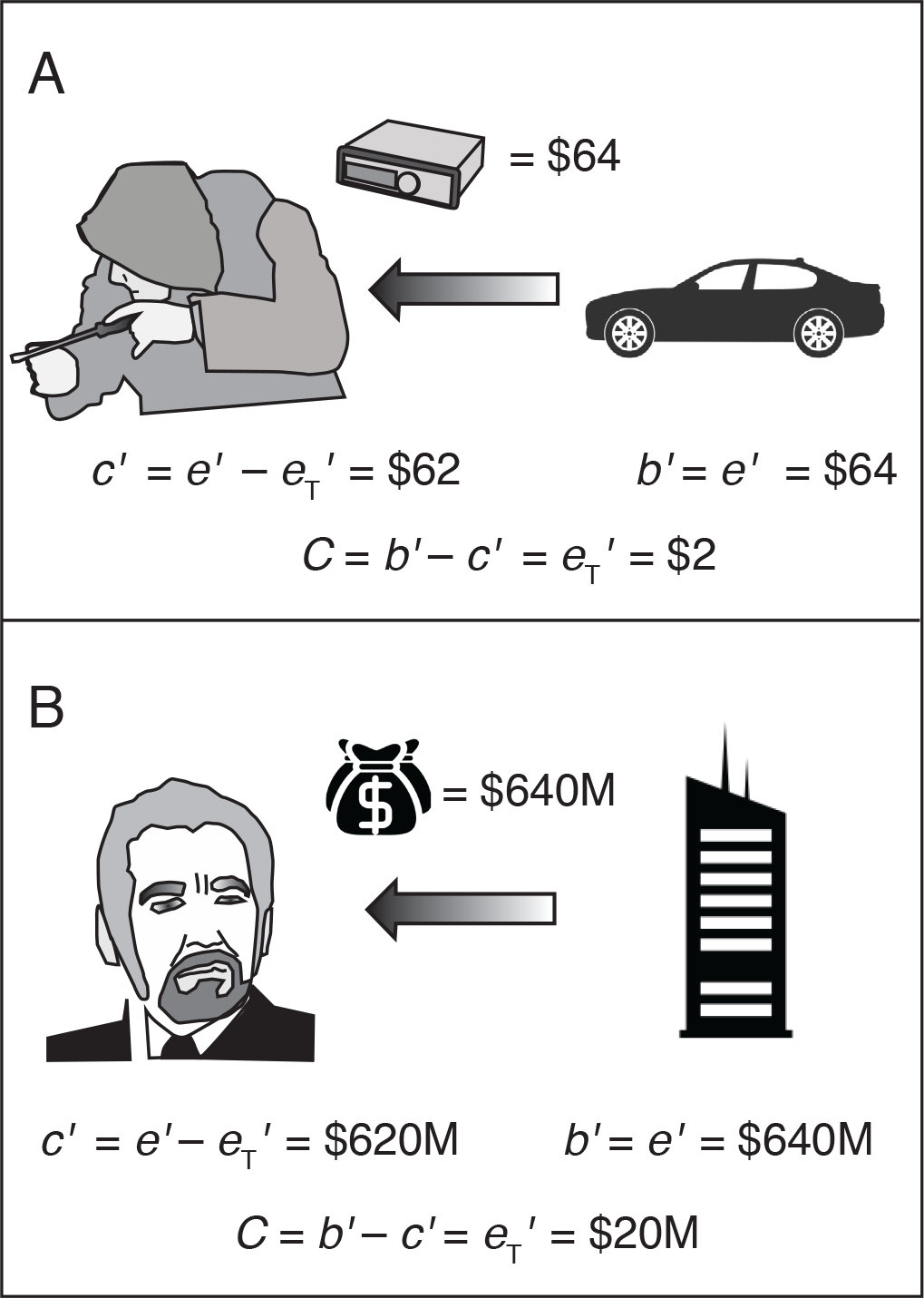
Why the net cost of conflict increases with the size of the resource at stake. *A*, A petty crook justifies spending $2 on a screwdriver to steal a car stereo worth $64. *B*, A high-level criminal justifies spending $20 million to steal bearer bonds worth $640 million. Because the net cost of conflict C increases with the energy spent in robbery 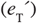, C tends to increase with the value of resource available for robbery. Moreover, a main requirement for harm to evolve is that the benefit to the actor is positive (*c′* > 0; Gilbert 2015), and this requires that the value of the resource stolen is larger than the effort of transferring the resource 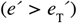. Here, for simplicity I here assume 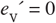.

From this argument, it is obvious why that coop-eration tends to carry a net benefit and conflict a net cost. It justifies our intuition for why, on average, we do best by associating preferentially with people with whom we will cooperate rather than conflict. Moreover, it suggests ways one might go about measuring the terms of Hamilton’s rule by a focus on the energy-transfer terms from which *b* and *c* arise (see below).

### Model

I note that a specialized association-theory model has already been developed for application to fusion-rejection association preference systems (Gilbert 2015). The primary goal here is to generalize the conceptual framework that underpinned formulation of this model. In appendix A, available online, I explain how Gilbert’s (2015) terms relate to the general association-theory terms used here (table 1), and I review his assumptions (fig. A1 and caption). My conceptual model here will make reference to Gilbert’s (2015) results where a rigorous model is necessary to support general concepts.

## The Steps

Based on the foundations thus provided, I outline a common historical process that characterizes major transitions in social evolution (summarized in table 3), described in detail below.

**Table. 3:** A summary of steps involved in the evolution of association preference during a major (or minor) transition.

| Steps | Preconditions | Evolutionary consequences (elements afterward taken as preconditions) | Reference |
| --- | --- | --- | --- |
| **1. Non association → Indiscriminate Association** | Conditions favoring adaptive a or incidental association, including a net automatic benefit of being in close proximity with conspecifics, selection to utilize a common resource, avoid predators, or other physical causes | **Indiscriminate association**, resulting in widespread association (aggregation or fusion) and, oftentimes, variability in relatedness between associates (aggregation or fusion partners). | Allee 1931<br>Buss 1982<br>Krause and Ruxton 2002<br>Aanen et al. 2008<br>Gilbert 2015<br>Ward and Webster 2016 |
| **2. Indiscriminate Association → Differential treatment** | Variability of relatedness between associates and opportunities for favoring kin or exploiting nonkin, allowed by close proximity with conspecifics, and the presence of cues indicative of relatedness | **Differential treatment** based on cues correlated with relatedness, for the benefits of favoring kin as predicted by Hamilton's rule, and for selfishly harming and cheating nonkin, as predicted by a separate "facultative exploitation" condition | Hamilton 1964 Fletcher and Michener 1987 Grosberg and Quinn 1989 Rousset and Roze 2007<br>Cornwallis et al. 2009<br>Gilbert 2015<br>fig. 5 |
| **3. Differential treatment → Treatment-cue association preference** | Treatment-cue-dependent effects of association, due to conflict with those differing in cues and cooperation with those sharing cues | **Treatment-cue association preference**, favored for the benefits of avoiding nonkin who differ in cues without avoiding nonkin who share cues | Gilbert 2015 |
| **4. Treatment-cue association preference → Dedicated-cue association preference** | Residual conflict with nonkin differing in treatment cues, despite selection to use treatment cues for association preference, because treatment cues are only somewhat useful for association preference | **Dedicated-cue association preference**, favored where the benefit of avoiding residual conflicts with nonkin who differ in treatment cues outweighs the benefits of association with nonkin who share treatment cues, providing a net benefit to avoiding nonkin indiscriminately of the treatment cue per se; instead, based on a separate, dedicated cue used only for association preference (the "dedicated" cue) | Gilbert 2015<br>fig. 6<br>table 4 |
| **5. Dedicated-cue association preference → Genetic kin recognition** | Advantage to rare alleles of a dedicated association preference cue, for avoiding associations with random individuals indiscriminately of treatment cue type per se | **Genetic kin recognition**, the accumulation of dedicated association preference cue polymorphism restricts associations to close kin | Gilbert 2015<br>fig. 7A |
| **6. Incidental effects of association preference on major transitions** | Treatment-cue association preference or dedicated-cue association preference | **Facilitation of transitions**, by controlling the spread of interspecific social parasites (either form of association preference) or promoting the evolution of intraspecific altruism (dedicated-cue only) | Buss 1982, 1987<br>Aanen et al. 2008<br>Bourke 2011<br>Gilbert 2015<br>fig. 7B<br>table A2 |

**Note:** In cases where treatment cues are not at all useful for association preference (table 4), evolution may proceed directly from differential treatment to dedicated-cue association preference.

### 1. Non Association → Indiscriminate Association

If an association is the context for a social action, then associations must evolve before social actions. Associations can originate in three main ways. First, association can evolve even where the presence of conspecifics may not be beneficial, for example where conspecifics must gather in an area due to the presence of a necessary resource like water, food, or shelter (Allee 1931; Alexander 1974; Krause and Ruxton 2002; Ward and Webster 2016). Here, associations may form without kinship, and the presences of conspecifics may be detrimental. Second, association can evolve as a “one-way” behavior where only one individual must express aggregation or fusion for associations to form. Here, association can evolve despite a net automatic cost, and even among nonkin (Hamilton 1971; Krause and Ruxton 2002).

Third, association can evolve where the automatic benefits outweigh the automatic costs (Buss 1982; Krause and Ruxton 2002; Aanen et al. 2008; Korb and Heinze 2016). With net automatic benefits, association can evolve as a one-way behavior among nonkin. Where two individuals must express the aggregation or fusion behavior for associations to form, however, individuals must find association partners. Thus, kinship may allow the initial local concentration of an allele for association (Gilbert 2015). However, kinship is not necessary for maintenance, since random individuals will also share the allele once it is common (Gilbert 2015). This holds even where the association behavior is costly because of the exclusivity of two-way association. For example, two social amoebae clones must both express cell adhesion genes *csA* to aggregate, and thus csA-bearing amoebae associate only with other csA-bearing amoebae (Ponte et al. 1998). Likewise, two *B. schlosseri* colonies must both express *fester* to fuse (Nyholm et al. 2006), and thus *fester*-bearing *B. schlosseri* colonies will fuse only with other *fester*-bearing colonies. Consequently, costly association behaviors can be stable among nonkin (Garcia and De Monte 2012).

It is important, however, not to confuse “association” and “altruism,” because altruism, in contrast to association, often requires kinship to be stable. For example, two social amoebae clones must both express cell adhesion genes *csA* to be included in aggregates, but they still may be exploited by *fbxA-* mutants (Gilbert et al. 2007). Most examples of the so-called “greenbeard altruism” effect (*e.g.*, Queller et al. 2003; Smukalla et al. 2008; Gruenheit et al. 2017) confuse “association” and “altruism.” In fact, although many organisms indiscriminately aggregate or fuse with nonkin (Krause and Ruxton 2002; Gilbert 2015; Ward and Webster 2016), there are few examples of altruism and division of reproductive labor among nonkin (Bourke 2011). Moreover, just because two clones aggregate or fuse, does not necessarily mean they will cooperate. As we all know from experience, “association” does not imply “cooperation.”

### 2. Indiscriminate Association → Differential Treatment

If individuals indiscriminately associate with conspecifics, relatedness among associates will depend on passive population structure. In the special case of very high relatedness or very low relatedness, only indiscriminate social actions are expected to evolve. Where relatedness among encountered individuals is intermediate or variable, due either to fusion or aggregation of non-clones, discriminatory social actions may evolve. Both discriminatory helpful and discriminatory harmful behaviors can be favored relative to indiscriminate actions or obligate non-actions given two conditions are met: (i) the discriminating nepotism condition, or Hamilton’s rule with *r* determined in part by cue variability; and (ii) the facultative exploitation condition, which specifies the advantage of exploiting nonkin (fig. 5). Briefly, the “discriminating nepotism” condition specifies the advantage of preferentially helping or avoiding harming kin. In the situation Gilbert (2015) considered, the discriminating nepotism is given by *rb* > *c* or *r b′* > *c′*, respectively, where *r* = *K* / [*K* + (1 − *K*)*P*_*i*_], *K* is the passive kin encounter rate, and *P*_*i*_ is the probability of randomly sharing the cue with a non-relative. Here, *r* is dependent on the passive kin encounter rate and variability of a cue used for differential treatment.

**Figure 5:**
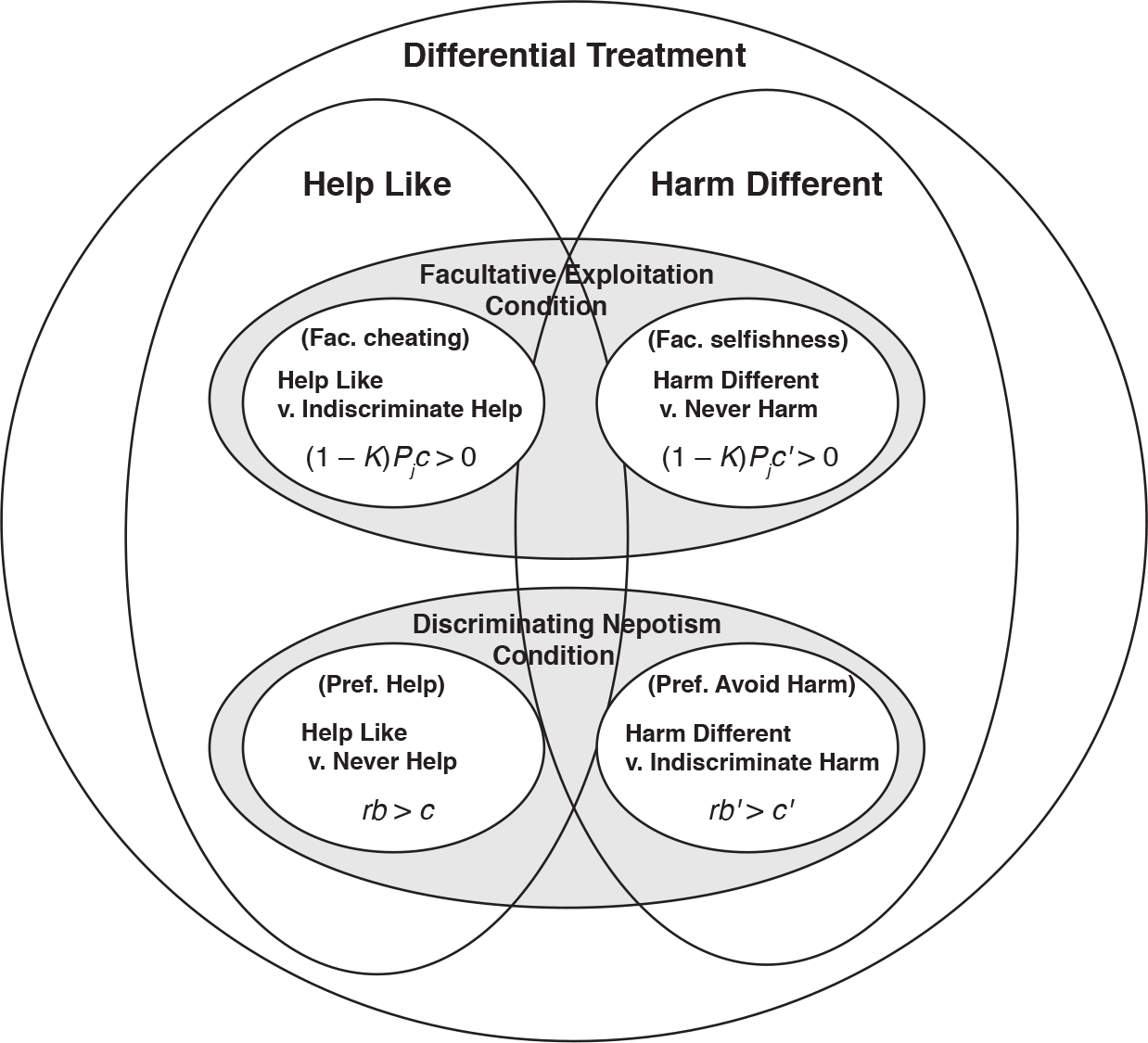
The evolution of differential treatment. The Venn diagram shows the two basic conditions required for the evolutionary stability of two forms of differential treatment, discriminatory social actions that help those sharing cues (“Help Like”) and harm those differing in cues (“Harm Different”). For both Help Like and Harm Different to be ESSs, the “facultative exploitat ion” condition and the “discriminating nepotism” condition (Hamilton’s rule for discrimination) must hold. The facultative exploitation condition manifests as either a cheating condition, in the case of help, or a selfishness condition, in the case of harm. Likewise, the discriminating nepotism condition manifests as a preferential helping of kin, or preferential avoiding harm to kin condition, in the case of Help Like and Harm Different, respectively. Here, *r* = *K* / [*K* + (1 – *K*)*P*_*i*_], where *K* is the passive clonemate encounter rate, and *P*_*i*_ is the probability of randomly sharing the cue with nonkin (table A2). See text for definition of terms.

The second condition for differential treatment is what I call the “facultative exploitation” condition. In the situation Gilbert (2015) considered, the facultative exploitation condition is given by, (1 − *K*) *P*_*j*_ *c* > 0 or (1 – *K*) *P*_*j*_ *c* > 0, respectively, where 1 − *K* is the probability of encountering nonkin, and *P*_*j*_ is the probability of not sharing a cue with nonkin. The former instance of the facultative exploitation represents cheating of altruism where *c* > 0, while the latter represents pure selfishness where *c′* > 0 (fig. 5). The importance of the facultative exploitation condition, relative to the discriminating nepotism condition, is that an organism might initially evolve indiscriminate altruism due to a passive population structure that restricts encounters to kin (e.g., a bottlenecked life cycle; Bourke 2011). However, if such an organism thereafter evolves to fuse (Buss 1982), the drop in relatedness can cause indiscriminate altruism to be invaded by discriminatory altruism, and obligate non-harm to be invaded by discriminatory harm. Both behaviors evolve according to the facultative exploitation condition (fig. 5).

Discriminatory helpful and harmful social actions, when fixed, select against polymorphism of their cues (Crozier 1986; Gilbert 2015). Thus, polymorphism must be maintained by extrinsic balancing selection (Rousset and Roze 2007).

### 3. Differential Treatment → Treatment-Cue Association Preference

If a cue is used for differential treatment, associating preferentially based on this cue allows an organism to avoid conflict while still entering beneficial or cooperative associations, including those with nonkin. Selection for such “treatment-cue” association preference will be strongest where the greatest resources are at stake, because the greatest net costs of conflict are likely to be found (fig. 4). For association preference to be favored, however, it is important that the cost of conflict outweigh the net automatic benefit, because this makes associations with random individuals costly enough to favor discriminatory “avoidance” (rejection or segregation) strategies over indiscriminate association (never reject or never segregate) strategies. Treatment-cue association preference will also most likely be the first behavior to evolve, even where segregation or rejection based on the cue does not completely avoid associations (table 4), because it is already used as part of a behavioral discrimination system (differential treatment). Treatment-cue association preference can lift a selective pressure imposed by discriminatory conflict against polymorphism, but it does not select for polymorphism (Gilbert 2015).

**Table. 4:** Examples of treatment cues classified according to their utility for association preference, and dedicated association preference cues.

| Treatment cue utility | Class | Example | Dedicated cue | Reference |
| --- | --- | --- | --- | --- |
| Not useful for association preference ($\delta = 1$) | Spatial locations | Social insects use location of nest as cue for help or harm | Spatial location cannot be used as a cue to discriminately reject foreigners at one location, so a separate phenotypic cue (i.e., odor) must be used | Fletcher and Michener 1987<br>Ratnieks 1991<br>Gamboa 2004<br>Gilbert 2015<br>fig. 7B |
| Somewhat useful for association preference ($1 > \delta > 0$) | Individual phenotypes detectable only in close proximity | Bats and birds use odor and facial patterning for differential treatment in close proximity | Odors and facial patterns are difficult to detect at a distance, so bats and birds use auditory cues to relocate offspring from a distance | Stoddard and Beecher 1983<br>Beecher 1988<br>Loughry and McCracken 1991<br>Balcombe and McCracken 1992<br>Strickler 2013 |
|  | Individual phenotypes detectable only at late stages of social development | Cues for facultative exploitation detectable immediately after slug formation, as suggested by reduced migration distances of chimeric slugs and slightly altered stalk:spore proportions | Social amoebae segregate immediately before slug formation based on cell adhesion genes | Foster et al. 2002<br>Buttery et al. 2009<br>Hirose et al. 2011<br>Jack et al. 2015 |
|  | Individual phenotypes detectable only at late stages of fusion | Cues for germline parasitism detectable only after fusion of blood vessels, hyphae, or plasmodia | Rejection based on cues detectable prior to substantial fusion | Smith et al. 2006<br>Rosengarten and Nicotra 2011<br>Clark and Haskins 2012<br>fig. 1B<br>fig. 7A |
| Perfectly useful for association preference ($\delta = 0$) | Individual phenotypes detectable before association | *B. schlosseri* overgrow or otherwise harm colonies that are *fuhc*-incompatible. *B. schlosseri* larvae preferentially settle near those that sharing *fuhc* itself | N/A | Grosberg and Quinn 1986<br>Buss 1990<br>Ben-Shlomo et al. 2008<br>Carpenter et al. 2011 |

**Note:** The parameter $\delta$ indicates the amount of an association that occurs despite activation of treatment-cue based avoidance. Thus, higher values mean the treatment cue is not useful for association preference.

### 4. Treatment-Cue Association Preference → Dedicated-Cue Association Preference

Consider a case in which a trait *A* (e.g., an odor), is detectable in close proximity and used for differential treatment (e.g., Loughry and McCracken 1991). It could be that a separate trait *B*, such as a loud cry, is more detectable at a distance (e.g., Balcombe and McCracken 1992). In that case, *B* would be more useful for avoiding associations before social actions are expressed contingent on *A*. More generally, if a cue *A* is detectable only at late stages of aggregation or fusion, it might initially be used as a cue for differential treatment. However, a separate cue *B*, detectable earlier during aggregation or fusion, may be more useful for discriminatory segregation or rejection (association preference). Thus, a change in the environment that results in the origin of a new conspicuous feature *B*, or the origin of some variability of a cue *B*, may allow the evolution of “dedicated-cue” association preference behavior based on a cue *B* that is detectable before social actions expressed on *A*.

Thus it may be expected that organisms that initially use treatment cues for association preference later evolve to use dedicated cues. For example, colonial ascidians may have initially used cues detectable in the blood to activate germline parasitism behavior, and only later seized upon a trait detectable on the outer tunic, *fuhc*, for use as a rejection cue (fig. 6A; Gilbert 2015). Alternatively, in some cases, only a dedicated cue can be used. For example, social insects use spatial cues for nest robbery, but spatial cues cannot be used to distinguish between individuals at one location. Thus, separate, phenotypic cues must be used for rejecting nonkin from nests (fig. 6B). In general, where treatment cues are not very useful for association preference (table 4), the use of separate “dedicated” cues may thus be favored.

**Figure 6:**
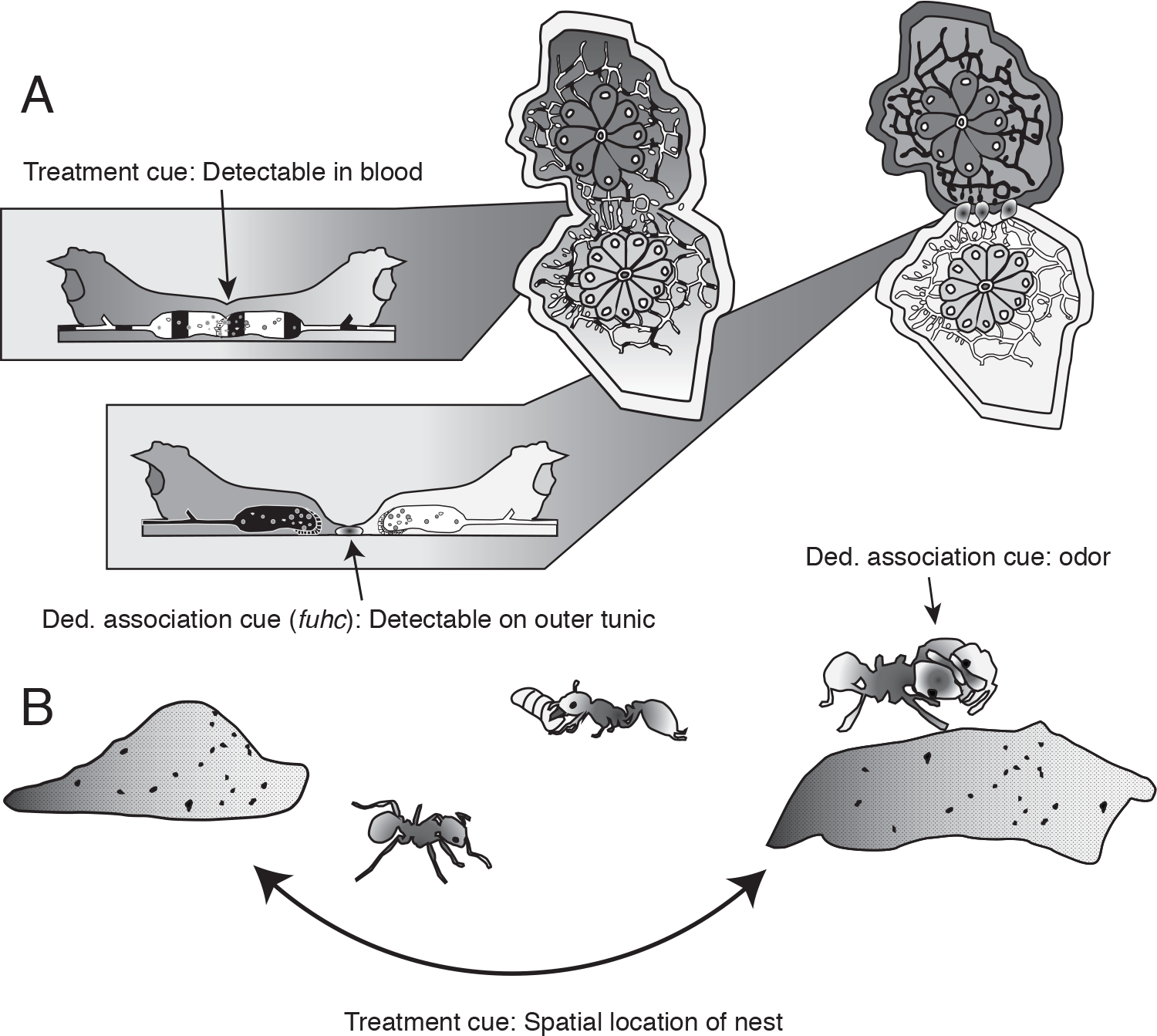
Examples of treatment cues not useful for association preference. *A*, Marine invertebrates *B*. schlosseri (Tunicata) may use phenotypic cues detectable after blood vessel fusion to activate germline parasitism, resulting in decreased growth rates and germline parasitism in chimeras (Rinkevich and Weissman 1989, 1992; Stoner and Weissman 1996; Laird et al. 2005). These are the putative “treatment” cues, and are postulated to be used for both differential treatment (germline parasitism) and treatment-cue association preference (delayed rejection, which occurs weeks after fusion of blood vessels [Rinkevich and Weissman 1989; Saito et al. 1994]). Rejecting on a separate, outer-more cue encoded by the fuhc is thus useful because it allows colonies to avoid the costs of fusion that occur with delayed rejection (Fig. 1Bi). Lateralview picture modeled after Taneda et al. (1985). *B*, Fire ant colonies Solenopsis invicta exhibit brood raiding behavior, selfishly robbing other colonies’ larvae. Brood raiding is more prevalent at times of year before colony recognition systems become functional (Balas and Adams 1996), suggesting spatial locations are used as cues for intraspecific parasitism, rather than colony odor cues. However, spatial locations cannot be used to recognize and reject foreigners from nests, so ants must use separate, phenotypic cues. Thus, in both marine invertebrates and fire ants, a separate cue can be used for association preference as for differential treatment.

What are the conditions necessary for the evolution of “dedicated-cue” association preference? In terms of balancing different forms of discrimination error (Sherman et al. 1997), dedicated-cue association preference will be favored when the cost of increased avoidance error, manifesting as the incorrect avoidance of those sharing treatment cues, is outweighed by the benefit of avoiding those who differ in treatment cues, allowed by using a more effective “dedicated” association preference cue. For example, where there are three treatment cue alleles at equal frequency, the chance of sharing the treatment cue randomly (with nonkin) is 1/3, while the chance of differing in the treatment cue randomly is 2/3 (fig. 7A). Dedicated-cue association preference will be favored when the cost of increased avoidance error, manifesting as the incorrect avoidance of the 1/3 of nonkin sharing treatment cues, is outweighed by the benefit of avoiding the 2/3 of nonkin who differ in treatment cues.

**Figure 7:**
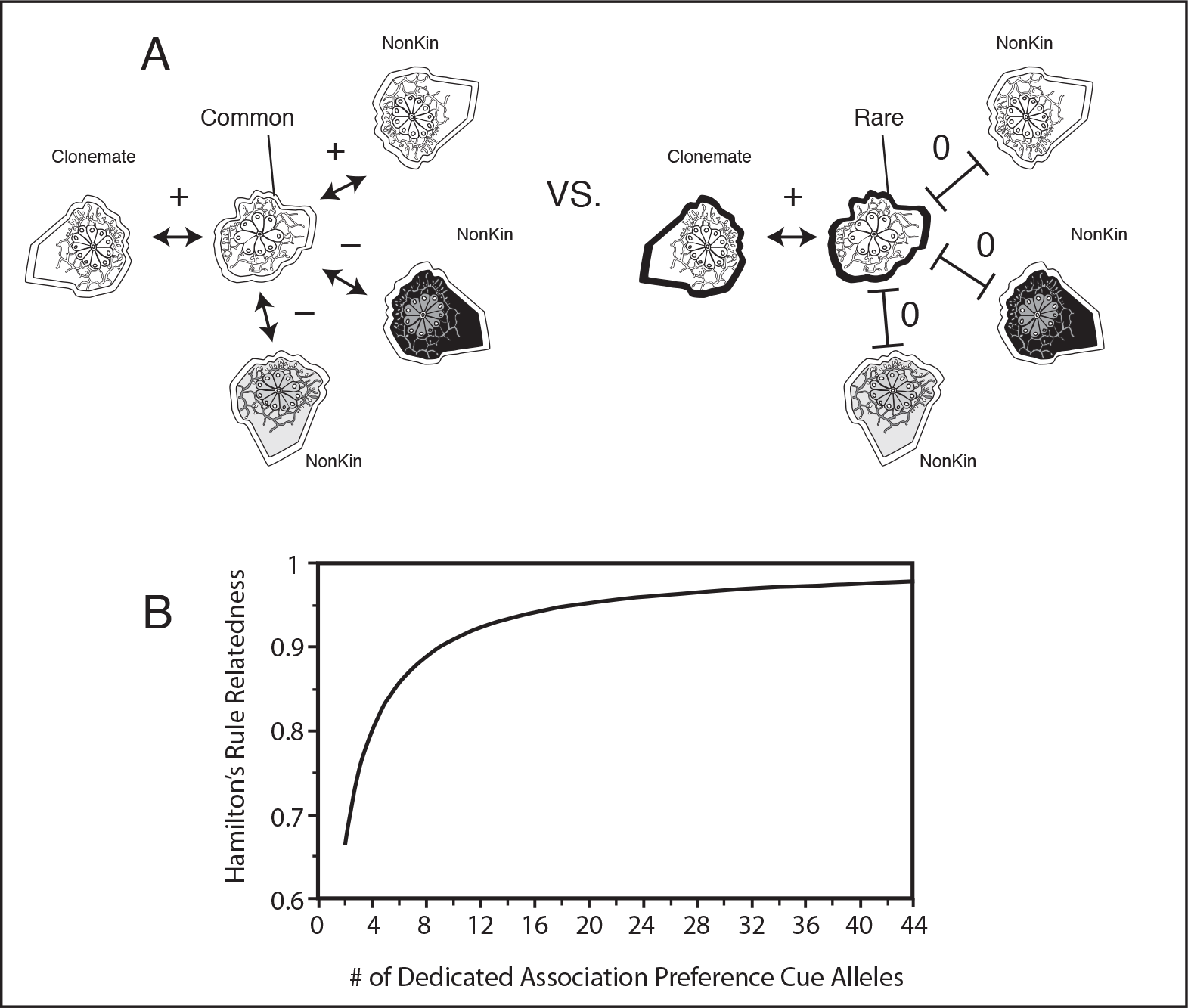
The evolution of genetic kin recognition. *A*, The reason polymorphism accumulates at the dedicated association preference cue locus can be understood by comparing the effect of a common (left) or rare (right) cue allele on organismal reproductive success. Individuals with common cue alleles will fuse with nonkin. Where the treatment cue gene has 3 alleles at equal frequency, nonkin will have 2/3 chance of conflicting, resulting in a negative effect of the association (fusion) with nonkin on average (where the cost of conflict outweighs the automatic benefit of association). Individuals with rare cue alleles, in contrast, will almost never fuse with nonkin, but still fuse with close kin. In the present discussion, it is assumed that the only kin are clonemates. B, The effect of increasing dedicated association preference cue polymorphism on the relatedness (r) term of Hamilton’s rule, for the conditions pictured in (A) in a haploid organisms that encounters either clonemates or nonkin. Specifically, it is assumed the treatment cue has 3 alleles at equal frequency, that the probability of encountering kin is *K* = .25, and nonkin is 1 − *K* = .75, in a randomly-mating population. Here, *r* = *K* / [*K* + (1 − *K*)*P*_*ii*_], where *P*_*ii*_ = *P*_*i.*_ * *P*_*.i*_ is the probability of randomly sharing both the dedicated association preference cue (*P*_*i*._) and differential treatment cue (*P*_*i*._) (See table A2 for payoff matrix used to derive r).

### 5. Dedicated-Cue Association Preference → Genetic Kin Recognition

Once present, dedicated-cue association preference exerts a selective pressure for rare alleles of its cue locus. The effect of possessing a common dedicated cue is to increase the fraction of the population with which an organism can associate (fig. 7*A*, left), while the effect of possessing a rare cue allele is to restrict the fraction of a population with which an individual can associate (fig. 7*A*, right). Because avoiding random associations must be favorable for the dedicated-cue association preference behavior to evolve, rare alleles of its cue locus will be favored where the behavior evolves adaptively. This holds even in the example where the treatment cue has only three alleles, as in the example above (fig. 7). Polymorphism is, moreover, expected to accumulate, because the build up of polymorphism at the dedicated-cue locus increases the stability of the dedicated-cue association preference strategy versus plausible alternatives (Gilbert 2015).

### 6. Long-Term Effects

Association preference can have two long-term effects on sociality. First, it can prevent the spread of reproductively-isolated, interspecific social parasites. Examples of social parasites include transmissible cancers in vertebrates and marine invertebrates, parasitic nuclei in red algae, and dedicated social parasites in vertebrates and arthropods (table A3). Both treatment-cue and dedicated-cue association preference can incidentally defend against interspecific social parasitism.

Second, the evolution of dedicated-cue association preference can increase relatedness relevant to the evolution of intraspecific altruism (fig. 7*B*). Dedicated-cue association preference has this effect because it avoids associations with nonkin who share the treatment cue, which disfavors intraspecific obligate cheaters relative to discriminatory altruists in the long term (table A2). Moreover, relatedness required for discriminatory harm is *r* > *c′* / *b′*, where 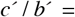 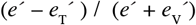; while relatedness required for discriminatory help is *r* > *c* / b, where *c* / *b* = (*e* + *e*_T_) / (*e* + *e*_V_). Thus, where energy-transfer terms are equal 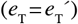, it is possible for discriminatory harm to evolve first (because *c* / *b* > *c′* / *b′*), select for association preference, and only later discriminatory altruism to evolve (Gilbert 2015).

## Measurements

### Basic Approach

Here, I will review basic approaches to measuring association-theory parameters. Consider first the simplest case of a species that forms association by aggregation or fusion, but which does not have differential treatment or association preference. The average baseline reproductive success of solitary individuals who never associate is *W*_0_. In turn, *W*_0_ + *ɛ* is the average reproductive success of individuals who associate with conspecifics, where *ɛ* is the net automatic benefit of association (automatic costs subtracted from automatic benefits). Thus, *ɛ* can easily be estimated as the difference in reproductive success of solitary and associated individuals.

Now, consider a case in which differential treatment is fixed based on a single “treatment” cue. Individuals who associate will have an average reproductive success of *W*_0_ + *ɛ* + *B* or *W*_0_ + *ɛ* – *C*, depending on whether the treatment cue is shared or not, respectively. If it were possible to experimentally inhibit the expression of social actions, then *W*_0_ + *ɛ* could be measured among associated individuals. This would allow an estimation of 8 as above, and in turn *C* and *B* could be measured by allowing social actions to be expressed as usual.

Measurements become more difficult once association preference is fixed. Where treatment-cue association preference is fixed based on a reliable cue (*δ* = 0), the various parameters can be measured in the same way as when only differential treatment is fixed (above). However, now individuals who differ in treatment cues avoid each other completely. Thus, it is impossible to measure the reproductive success of conflicting individuals *W*_0_ + *ɛ* – *C* unless individuals are experimentally forced to associate. This could be accomplished by silencing genes associated with association preference behavior, or using laboratory techniques that force association among those who would otherwise avoid each other. Mycologists often, for example, force chimerism between fungal strains differing in heterokaryon incompatibility genes (e.g., Cortesi et al. 2001). In some cases, however the parameter *δ*, indicating how useful a treatment cue is for association preference, may actually be a composite function of multiple cues. Studies of the Chestnut blight fungus *Cryphonectrica parasitica* show that five loci contribute to putative treatment-cue association preference (heterokaryon incompatibility; Cortesi et al. 2001). Where *δ* is a composite function, there is also an additional measurement problem of dealing with multiple treatment-cue genes.

The greatest challenge in measuring the association-theory parameters comes when association preference is fixed based on a “dedicated” cue. Now, it is necessary to silence association preference based on both the treatment cues and dedicated association preference cues in order to disentangle *W*_0_, *ɛ*, *B*, and *C*. Moreover, separate questions arise. Presumably, where dedicated-cue association preference is fixed, the treatment cue is not perfectly reliable for association preference (*δ* > 0). Thus, another problem is in measuring the parameter *δ*. One way to measure *δ* is to study associations among individuals who share dedicated cues (e.g., major histocompatibility cues), but who differ in treatment cues (e.g., minor histocompatibility cues). In colonial ascidians *B. schlosseri*, for example, delayed rejection reactions based on “minor” histocompatibility genes take weeks to occur, in contrast to hours for the major histocompatibility cue (Rinkevich and Weissman 1988, 1989; Saito et al 1994). This suggests that *δ* in these organisms is quite high, i.e., that treatment cues are detectable late in fusion.

### Testing Hamilton’s Rule

I now provide a link to former theories by explaining how to test Hamilton’s rule using an empirical example. To do so, I will first explain how Hamilton’s rule was previously applied to explain association. I will then review how that this application of Hamilton’s rule yields an inconsistency, which is removed by distinguishing “social action” and “association.” Finally, I will explain how association theory can be used to test Hamilton’s rule.

Consider the textbook application of Hamilton’s rule to explain pairwise lek formation in male turkeys (Davies et al. 2015). Here, *r b* – *c* > 0 was formulated as follows. *r* was calculated as the mean pairwise relatedness between associates (lekmates), *b* as the mean difference in reproductive success between dominants and solitary individuals, and *c* as the mean difference in reproductive success between subordinates and solitary individuals, yielding *r b* – *c* = +1.7 (Krakauer 2005). This appears to be a successful testing of Hamilton’s rule, but it yields an inconsistency. In a similar example of association, unrelated ant foundresses form pairwise coalitions during colony founding, but only one foundress ultimate benefits by becoming “queen,” while the other is killed (Bernasconi and Strassmann 1999). If one applies the same methodology to measuring Hamilton’s rule for altruism, it must fail because *r* = 0.

Association theory, in contrast, predicts that both turkeys and ant foundresses form associations initially due to automatic benefits, and that associations are maintained because: (i) after conflict evolved, the automatic benefits of association outweighed the net costs of conflict, (*ii*) individuals do not know whether they will become dominants or subordinates prior to the establishment of associations, and (iii) subordinates would suffer higher costs by leaving or otherwise cannot leave. In appendix A, I provide evidence for these predictions.

How do we test Hamilton’s rule correctly? Association theory predicts that Hamilton’s rule for restraint on harm is not met in both cases. As applied to the unrelated ant-foundress example, for *r b′* < *c′* to hold, it is necessary only that *c′* > 0 (because *r* = 0). This is known because one foundress benefits by killing her partner and becoming the sole queen (recall, *c′* is the benefit of harm, or cost of harm avoidance, to the actor). In the turkey example, however, the main prediction is that turkeys harm lekmates related by *r* on average because the cost of harm-avoidance to the actor is not greater than *r* times the benefit of harm-avoidance to the recipient. How can we test Hamilton’s rule in this latter case?

To test Hamilton’s rule, I assume that without any conflict, each associated turkey has on average a reproductive success of *W*_0_ + *ɛ*. By being in continual close proximity, however, a dominant turkey can channel the reproductive success of its partner, equal to *e′* = *W*_0_ + *ɛ*, by usurping all of the copul tions with females. Here, I assume 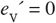, because each turkey would benefit equally from mating with females (contrasted to an asymmetric situation; fig. 3*B*). I assume, however, 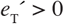 because there is energy spent in fighting. Because *b′* = *e′* and 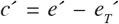, Hamilton’s rule *r b′* – *c′* > 0 becomes

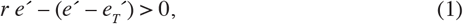

where *r* = .42, *e′* = 1/2 (*W*_0_ + *ɛ*) and 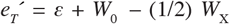. According to Krakauer 2005 measurements,*W*_0_ = .93 and *W*_*X*_ = 6.125, and therefore association theory predicts *ɛ* < 3.38 (see appendix A for details). This prediction can be tested by measuring *ɛ* among turkeys that do not suffer conflict, as may be allowed by experimentally preventing aggressive behavior.

## Discussion

Is the distinction between a social action and an association merely specialized and technical, or is the distinction fundamental to a theory of social evolution? A relatively im artial answer to this question requires appeal to data. In what follows, I will first explain the most general prediction of association theory, and then how association theory resolves the most outstanding anomaly of previous social theory. I will then discuss the insights of association theory for major transitions. I will finally focus on association theory’s assumptions, and end with an explanation of its most general theoretical implications.

### General Prediction

The most general prediction of association theory is that association preference evolves in response to differential treatment, most particularly in response to discriminatory harm. This prediction is more general than the prediction for cue polymorphism, because it can hold even where cue polymorphism does not evolve adaptively (i.e., it can hold for treatment-cue association preference). The prediction explains why association preference evolves in species lacking altruistic behavior (West-Eberhard 1989), and why fusion often results in net costs even though fusion was formerly perceived as “cooperation” (Aanen et al. 2008). According to association theory, association is not cooperation, but the context for a social action; and therefore association preference can evolve in response to discriminatory harm, which is why discriminatory harm and association preference cooccur (table 5).

**Table. 5:**
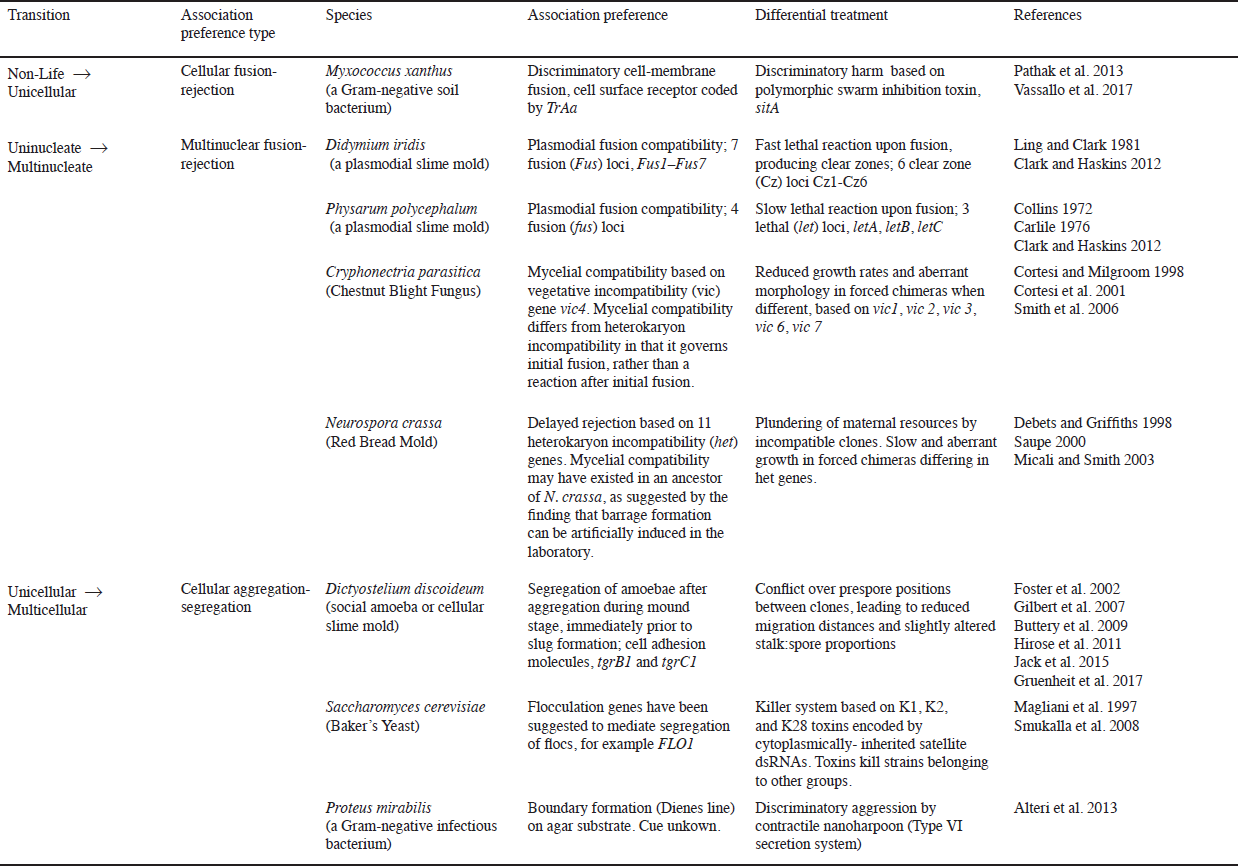
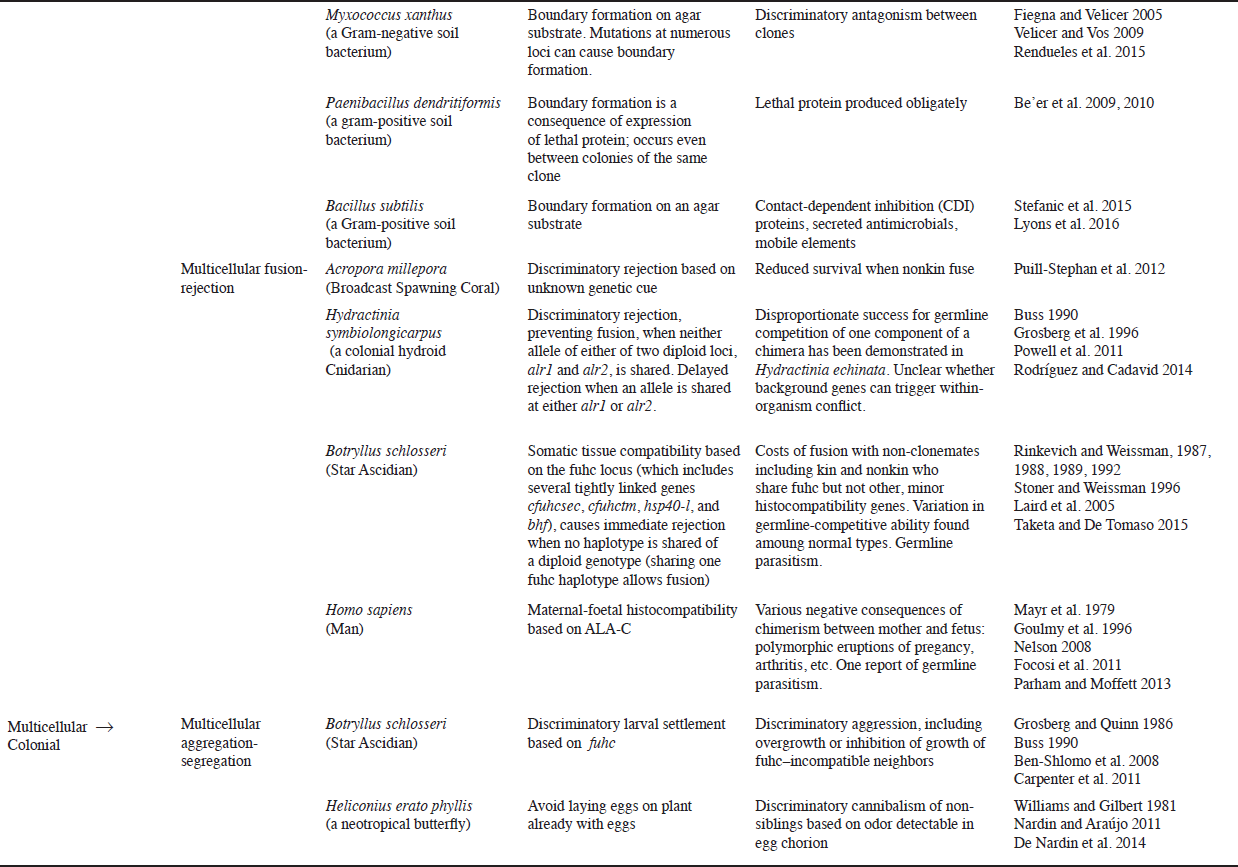
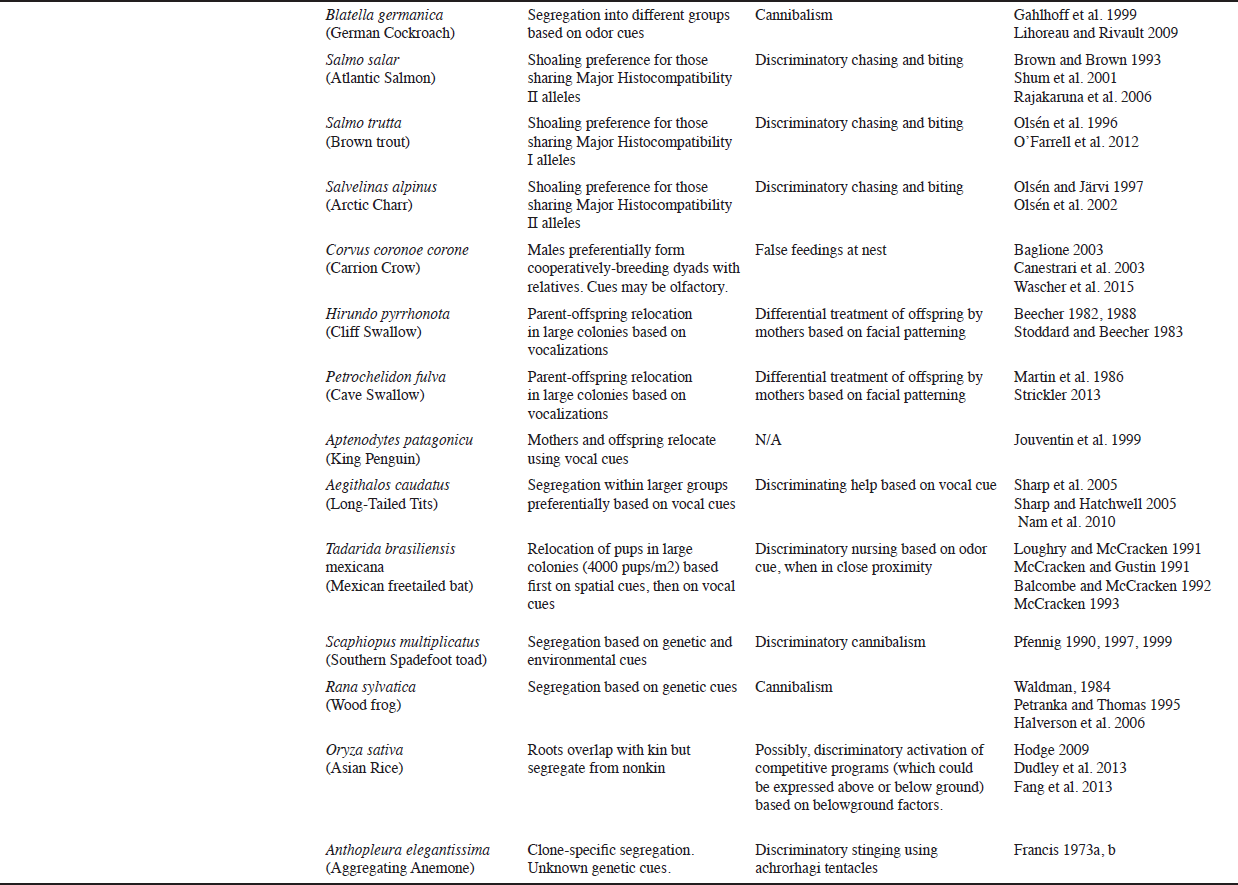
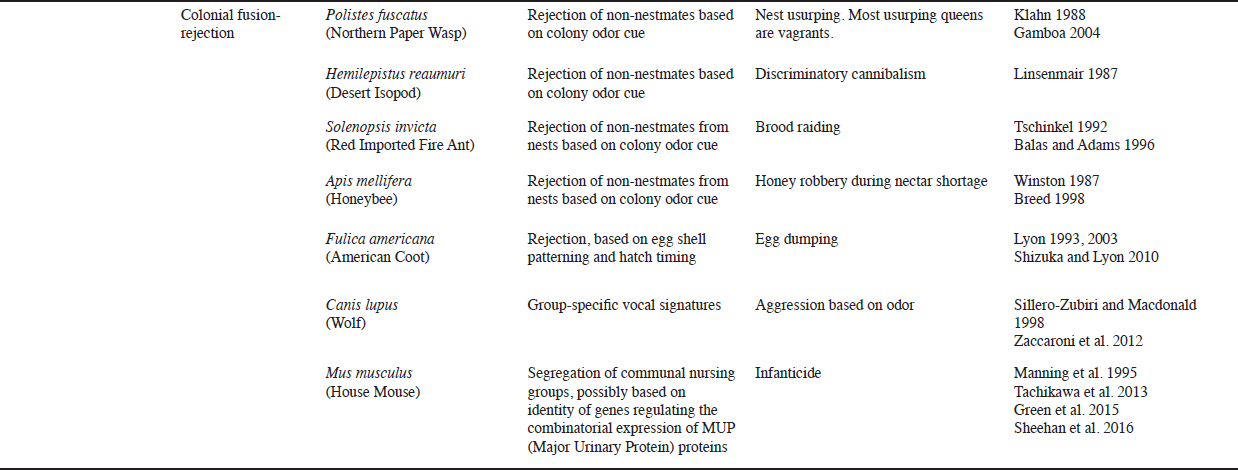
Association preference and differential treatment.

### Genetic Kin Recognition

The most famous anomaly of previous social theory, which directly resulted from confusing “social action” and “association,” is the anomaly of genetic kin recognition, also known as Crozier’s paradox (Crozier 1986; Aanen et al. 2008). Under a paradigm that confuses “social action” and “association,” authors typically applied models of discriminatory help or harm to explain the variability of cues used for association preference (Crozier 1986; Rousset and Roze 2007). When discriminatory help or harm are fixed, however, common cue alleles are favored for increasing the fraction of a population with which an individual can cooperate or avoid con-flict (Crozier 1986; Grosberg and Quinn 1989; Rousset and Roze 2007). It is therefore impossible for polymorphism to indefinitely accumulate because of selection imposed by differential treatment. The accumulation of some polymorphism allows the discriminatory behavior to fix, which then selects against polymorphism (Rousset and Roze 2007). What is necessary is a discriminatory behavior that, when fixed, favors polymorphism of its cue. Association theory resolves Crozier’s paradox by showing that polymorphism evolves for the purpose of association preference in response to differential treatment (particularly discriminatory harm), rather than for the purpose of differential treatment itself. This distinction would not have been possible under the previous definition of “kin recognition” as “differential treatment (Sherman et al. 1997)”.

The resolution of Crozier’s paradox yields association theory’s most specific prediction. Where a cue is used for association preference and there is no evidence of extrinsic balancing selection maintaining cue polymorphism, association theory predicts a separate genetic cue triggers discriminatory harm (fig. 7). Recent studies of *Myxococcus xanthus* bacteria confirm this prediction. In *M. xanthus*, there is no obvious extrinsic selective pressure for polymorphism of a discriminatory rejection cue, *TraA*, used for cell-level fusion-rejection association preference (Pathak et al. 2013). Consistent with the polymorphism evolving for the purpose of association preference in response to discriminatory harm, a separate cue was found that triggers harmful toxin production, a variable lipoprotein-coding gene *sitA* (Vasallo et al. 2017). It is important to note *TraA* is an “association” cue not a “help” cue.

From the hindsight of association theory, the solution to Crozier’s paradox seems obvious. All that is required is a distinction between forms of discriminatory behavior and their cues. However, a full solution also required a model of differential treatment that explains the relationship between discriminatory help and harm (fig. 5), and their consequences for net effects of interactions (fig. 2). It also required an explanation for why different cues would be used for association preference as for differential treatment (fig. 6 and table 4), and the relationship between variability of treatment cues and a dedicated association preference cue (fig. 7). Moreover, the solution requires explaining why association preference evolves where the greatest resources are at stake (fig. 4), and a logically-consistent explanation for all stages of social-evolutionary transitions including the origin of association (table 3). Finally, a solution was partially obscured by the narrow definition of kin recognition as “differential treatment (Sherman et al. 1997),” which in addition to conflating the discriminatory behavior and recognition ability, precludes a role for kin recognition in association preference.

### Major Transitions

Association theory suggests that effects of association preference on major transitions are largely incidental. Particularly, it suggests that a control on the spread of degenerate cheater mutants, or social parasites, is an incidental effect of adaptation to avoid discriminatory conflict. Indeed, differential treatment is found ubiquitously with all forms of fusion-rejection and aggregation-segregation association preference (table 5). In contrast, social parasites are restricted in their distribution. Where social parasites do occur, they have usually evolved to evade pre-existing recognition systems (e.g., social parasites in eusocial hymenoptera, and transmissible cancers in vertebrates; table A3). In certain birds, variability of egg shell patterns may have evolved in response to conspecific brood parasitism, and incidentally helped protect against interspecific brood parasitism (Samas et al. 2014; Lyon et al. 2015). In other cases, social parasites evolve in taxa that lack association preference systems (e.g., clams and red algae; table A3). Thus, association preference systems may evolve in response to differential treatment and only incidentally prevent the spread of social parasites.

The mere findings of association preference and genetic kin recognition are therefore not primae facie evidence of their importance for major transitions. Rather, hypotheses for long-term effects must be critically tested. For example, if it is postulated that genetic kin recognition maintains altruism, this hypothesis cannot be tested merely by showing that a genetic kin recognition exists (Hirose et al. 2011), that it is precise (Gruenheit et al. 2017), or that it might have an effect on maintaining altruism (Ho et al. 2013). Rather, experiments capable of disproving the hypothesis must be used (Gilbert et al. 2012). In cases where the hypothesis is refuted, alternative explanations for why altruism is maintained, for example based on passive population-structuring mechanisms, may be proposed (Gilbert et al. 2012; Smith et al. 2016).

Perhaps due to a lack of rigorous hypothesis tests for the importance of association preference for major transitions, many authors have assumed that a bottlenecked life cycle, or development by “staying together” of offspring, is sufficient for promoting the population structure required for a major transition (Maynard Smith and Szathmáry 1995, p. 8; Fisher et al. 2013, p. 1; Tarnita et al. 2013, p.20). Association theory suggests that a bottlenecked life cycle, though probably necessary, is not always sufficient (see also Buss 1982, 1987). Indeed, many organisms that go through a bottlenecked life cycle may fuse (fig. 2, table 5). Association theory brings renewed attention to importance of fusion-rejection systems in stabilizing major transitions (table 2), by showing how association preference can incidentally prevent the spread of socially-disruptive social parasites (table A3), and promote the evolution of altruism (fig. 7).

What is the evidence for long-term effects? One example where association preference may have stabilized a major transition is found in eusocial hymenoptera. In contrast to marine invertebrates that secondarily evolved to fuse (Cohen et al. 1998; Gilbert 2015), in social insects free mobility and frequent mixing of individuals from different colonies could have been a default before the evolution of altruism and eusociality. From a plausibility standpoint, nestmate recognition fulfills the requirement of “dedicated-cue” association preference, suggesting it could have both constrained the evolution of social parasites, and facilitated the evolution of intraspecific altruism (fig. 6*B*). Indeed, phenotypic-cue based nestmate recognition could increase relatedness between actors and recipients of spatial-cue based altruism. Consistent with an effect on the evolution of altruism, the loss of nestmate recognition systems can reduce relatedness within nests (Helanterra et al. 2009). Consistent with an effect on defending against social parasites, most social parasites in eusocial hymenoptera have evolved sophisticated mechanisms to evade nestmate recognition (Bourke 2011; table A3). Moreover, taxa that lost nestmate recognition have a “twiggy” phylogenetic distribution, suggesting they could have been extinguished by the spread of intraspecific obligate cheaters or interspecific social parasites (Helanterra et al. 2009).

### Assumptions

The most important assumption allowing the construction of a detailed model is that associations are symmetric. This assumption is justified on grounds that even organisms with complex sensory systems and brains (e.g., vertebrates) usually require some time in association, before it becomes apparent who is “stronger” in a contest (hence, ritualized tactics; Maynard Smith 1982). Thus, one might expect simpler organisms to also be incapable of gauging the outcomes of association before associating. Consistent with this, colonies of clonal anemones *Anthopleura elegantissima* of different competitive abilities coexist side by side in nature, suggesting they are unable to judge relative competitive ability (Ayre and Grosberg 1995). In clonal organisms that do not senesce, however, it is possible that alleles coding for strategies could be favored to behave as if they can gauge relative strength in conflict. For example, an allele associated over evolutionary time with a particularly “strong” genetic background might benefit by adopting a strategy that takes advantage of this fact. How might the results change if we allow for asymmetric association? Consider a strategy that avoids associations with individuals who are stronger and also differ in the treatment cue. Such a strategy could be favored for entering conflict with only weaker individuals. However, it could still be invaded by a dedicated-cue association preference behavior if the treatment cue is not obvious early enough (*δ* ≫ 0). Thus, it might be expected that association preference would often evolve anyway.

Another assumption allowing a simple model is that the only kin encountered are clonemates. This assumption does not detract seriously from a model in which association preference and genetic kin recognition evolve to avoid associations with nonkin (fig. 7). However, one question that may be answered only by considering encounters with sexual kin is why some marine invertebrates allow fusion with parents and siblings (Grosberg 1992). Association theory suggests that fusion with sexual kin could be favored because they are more likely to share treatment cues. Indeed, fusion with kin in *B. schlosseri* is, on average, less costly than fusion with nonkin in the laboratory (Rinkevich and Weissman 1992). If automatic benefits are greater in nature (Chadwick-Furman and Weissman 1995), fusion with sexual kin might carry a net benefit in nature. Moreover, the historical perspective of association theory suggests that in a case where polymorphism has already evolved based on a “complete-matching” mechanism that restricts fusion to clonemates, a mechanism that allows fusion with sexual kin (a “partial-matching” mechanism, Grosberg 1988, p. 398) would much less often allow fusion with nonkin, but still quite similarly allow fusion with kin. This could allow kin-fusion to evolve (see appendix A for discussion of a few more assumptions).

### A Dominant Framework?

The most dominant general framework today is “generalized Hamilton’s rule theory.” This paradigm applies Hamilton’s rule to explain traits causing the formation of groups (Queller 1985; Bourke 2011), maintenance of groups (Vehrencamp 1983; Nonacs and Hager 2011) and fusion of organisms (Buss and Green 1985; Aanen et al. 2008). Theories that use Hamilton’s rule to explain association decisions, like whether to stay in a group or leave (Vehrencamp 1983; Nonacs and Hager 2011), fall within the umbrella of this “general Hamilton’s rule theory.” Even supposed alternative paradigms, like Nowak et al.’s (2010) “theory of eusociality,” and Ryan et al.’s (2015) “social niche construction theory,” are part of the same paradigm, because they also extrapolate from models of social actions to explain association phenomena (Hamilton [1964] himself, however, applied Hamilton’s rule only to indiscriminate social actions). For example, both Nowak et al. (2010, p. S29) and Ryan et al. (2015, p. 64) explain traits affecting population structure (e.g. dispersal or aggregation) with models of altruism, by assuming the two different traits are linked and co-evolve (*sensu* Powers et al. 2011).

Association theory differs from the previous paradigm because it does not extrapolate from models of social actions to explain associations. Instead, it explains association in its own right. It draws on automatic effects of association to explain the initial origin of association, and it specifies the roles of automatic effects and net effects of social interactions in selecting for association preference and genetic kin recognition. Within association theory, Hamilton’s rule is subsumed as a narrow statement describing the benefit of nepotism, important for explaining the evolution of indiscriminate and discriminatory social actions. Hamilton’s rule alone, however, is not even sufficient to explain the stability of discriminatory social actions; the facultative exploitation condition is also necessary (fig. 4). Earlier models invented for the purpose of extending Hamilton’s rule (or other models of social actions) to cover association phenomena will probably be eventually disregarded or revamped for a new purpose under association theory.

## Conclusion

Association theory is a new framework for analyzing social evolution based on the core distinction between “social action” and “association.” Translating this simple distinction, which we all probably make every day of our lives, into a theory of social evolution yields subsidiary distinctions between “automatic effects of association” and “benefits of cooperation,” and between “differential treatment” and “association preference.” It also yields the following predictions: (*i*) aggregation and fusion normally evolve before cooperation, according to “automatic benefits” of association; (ii) social actions often evolve as “differential treatment” behaviors that favor kin and exploit nonkin; (*iii*) association preference typically evolves in response to differential treatment to avoid conflict with nonkin; (i*v*) where separate cues are used for association preference as for differential treatment, cue polymorphism can evolve adaptively for the purpose of association preference itself, yielding adaptive genetic kin recognition; (*v*) association preference can incidentally prevent the spread of interspecific social parasites or promote the evolution of intraspecific cooperation. These predictions are imminently testable.

## Acknowledgments

I thank L. Gilbert, C. Hawkes, and C. Bensimon for encouragement and support. I thank M. Ryan, M. Amcoff, and L. Gilbert for discussions. I thank J. Marshall, J. Andre, P. Taylor, Y. Stuart, and L. Gilbert for comments. I also appreciate the invitation from J. Strassmann and D. Queller to attend the 2015 “Organismality” conference in Washington University, St. Louis, MO, which was funded by the John Templeton Foundation. I thank attendees of the conference for discussions, including B. Swalla, R. Grosberg, N. Tsutsui, B. Johnson, E. Herre, j. smith, D. Haig, and D. Aanen. I express gratitude to Leo Buss, J. Strassmann, and D. Queller for inspiration leading to this work. I also appreciate the hospitality of the members of the Brackenridge Field Laboratory at the University of Texas Austin.

